# pytc: a python package for analysis of Isothermal Titration Calorimetry experiments

**DOI:** 10.1101/234682

**Authors:** Hiranmayi Duvvuri, Lucas C. Wheeler, Michael J. Harms

## Abstract

Here we describe *pytc*, an open-source Python-package for global fits of thermodynamic models to multiple Isothermal Titration Calorimetry experiments. Key features include simplicity, the ability to implement new thermodynamic models, a robust maximum likelihood fitter, a fast Bayesian Markov-Chain Monte Carlo sampler, rigorous implementation, extensive documentation, and full cross-platform compatibility. *pytc* can be used as either a programming API or with a GUI. It is available for download at: https://github.com/harmslab/pytc.

## Introduction

Isothermal Titration Calorimetry (ITC) is a powerful technique for measuring the thermodynamics of inter-molecular interactions (1,2). It provides information about the free energy, enthalpy, entropy, and stoichiometry of binding. Combining information from ITC experiments under different conditions can reveal further properties, such as heat capacity change or the number of protons gained or lost during binding (3–5).

Extracting thermodynamic information requires fitting models to observed heats, potentially across multiple experiments. Several software packages exist for fitting these models. These include commercial software that ships with instruments. These packages are closed-source and cannot be easily extended by users who wish to analyze different or more complex models. A powerful alternative to this approach is *SEDPHAT* (6) (and its recent fork *ITCsy* (7)). These packages allow for global fitting of exceedingly complex models (6, 8). While these packages are powerful, it remains difficult for users to extend these packages with new fitting models. They also require proprietary software: either Windows (SEDPHAT) or Matlab (ITCsy).

Here we report *pytc*, a flexible Python-based package for fitting ITC data. It is open source, released under the Unlicense. It is designed for ease of use for simple fitting scenarios, but also has ready extensibility and flexibility for more complex fitting. It has a graphical user interface (GUI) for basic fitting, as well as a straightforward object-oriented application programming interface (API). The API allows users to write new models and integrate with other analysis pipelines. The software depends on the python extensions numpy (9, 10), scipy (11), and matplotlib (12). The GUI also requires PyQT5.

- API download at https://github.com/harmslab/pytc (13)
- GUI-download: https://github.com/harmslab/pytc-gui (14)
- Tutorial and API examples as Jupyter notebooks: https://github.com/harmslab/pytc-demos (15)
- Software and model documentation: https://pytc.readthedocs.io/

## Package Description

The basic work flow is similar when using either the API or GUI. The user provides files that hold the heat-per-shot for each experiment, determined by integrating raw power curves using the instrument software or an alternative such as NITPIC (16). The user then specifies an individual fit model for each experimental data set. The current version of the software implements four such models: a blank titration, a single-site binding curve (1), an N-site binding polynomial (17), and a 1:1 competitor model (18).

These individual experiments can then be linked together into global fits. In the simplest case, one can link individual fit parameters to global parameters. For example, a user can fit a single binding constant and enthalpy to multiple experimental replicates. More complicated linkages can also be defined. The current release of the software implements three such linkages: proton-linked binding versus buffer ionization enthalpy (3), a Van’t Hoff analysis assuming a fixed enthalpy change, and a Van’t Hoff analysis assuming fixed heat capacity change (reviewed in (5)). For these models, individual experiments are done as a function of buffer ionization enthalpy or temperature, which then allows calculation of the enthalpy and binding constant of each individual experiment under its specific conditions.

### *pytc* is designed for rigorous model fitting

For maximum likelihood fits, it uses `scipy.optimize.least_squares` for regression (11). *pytc* also includes a Bayesian Markov-Chain Monte Carlo (MCMC) sampler, implemented using the *emcee* python package (19), that allows global Bayesian analyses of each model to be performed. After fitting/Bayesian sampling, the software returns extensive information useful for determining fit quality and performing model selection. This includes the fit RMSD, Bayesian credibility regions, number of fit parameters, log likelihood, the F test-statistic, and several variants of the Akaike Information test-statistic (20). Fit residuals are plotted for each fit, allowing selection of models with random residuals. Users can also access a corner plot that shows correlations between the fit parameter uncertainty distributions.

Another useful feature is fit reproducibility. When fits are done using the API, the exact steps taken are encoded as a standard python program. As a result, all fit guesses and manipulations are automatically recorded and the fits can be reproduced at will. This takes away the challenge of having to reproduce a series of steps done in a GUI. It also allows easy assessment of previously hard-to-quantify aspects of the fit, such as the sensitivity of the results to fit parameter guesses. The API is simple and does not require extensive programming background to use. This can be seen below, which shows a complete program for fitting a single-site binding model to a titration.

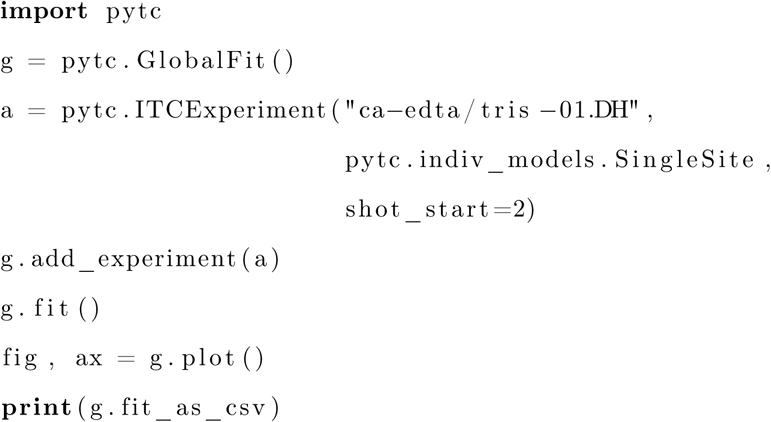

One of the key features of *pytc* is the ability to implement new binding models. This can be done for both models of individual ITC experiments and for global models. Implementing new models is straightforward and well documented. The coding required can be quite minimal. The single-site binding model included with the package is defined in 13 lines of python code. The global model linking buffer ionization enthalpy to proton gain/loss is defined in six lines. To illustrate the simplicity, the model is copied in full below:

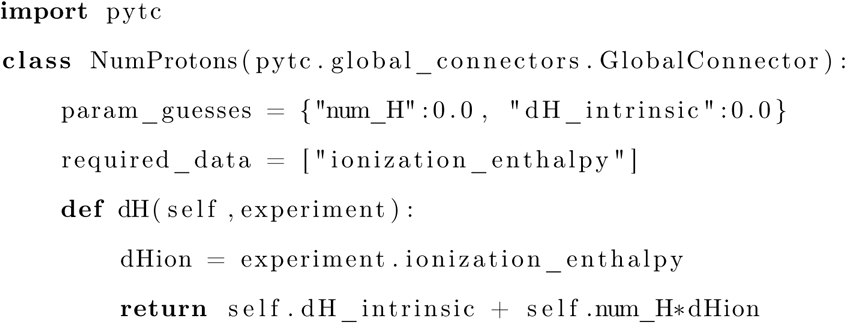

Importantly, the requirements for new models are specific and well-defined. A complete description of how to implement new models is available at https://pytc.readthedocs.io/en/latest/writing_new_models.html.

## Documentation

Finally, the software documentation contains detailed descriptions of how to use the software, details regarding the thermodynamic models, and notes about the statistical tools available in *pytc* (https://pytc.readthedocs.io/en/latest/). To help users gain traction with the software, we have created video tutorials for the GUI (https://pytcgui.readthedocs.io/en/latest/how_to_img.html) and have released demos with practice data for learning the API (https://github.com/harmslab/pytc-demos).

## Validation

To both validate and demonstrate the utility of *pytc*, we performed a series of fits to **ITC** binding experiments, ranging from simple single-site fits to global analyses involving multiple experiments. We demonstrate both the maximum likelihood and Bayesian MCMC approaches.

### Fitting single-site models

We first compared single-site fits using *pytc* to fits using the software that shipped with our VP-ITC (*Origin* 7.0552). We titrated 1.6 *mM CaCl*_2_ onto 0.1 *mM EDTA* in 100 *mM Tris*, *pH* 7.40 at 25 °*C* using a VP-ITC (GE Healthcare) (Figure 1A). All buffers were treated with CHELEX (≈ 1 *g* · *L*^−1^, stirring for 30 minutes), filtered at 0.22 *μm,* and then thoroughly degassed. Shot size was 1 *μL* for the first shot, followed by 5 *μL* for all following shots. Reference power was 7 *μcal* · *s*^−1^; stir speed was 633. For the *Origin* fit, we corrected for dilution by fitting a line to a blank titration—identical to the production titration except for having no EDTA—and then subtracting this line from the production heats. We discarded the first two shots before fitting. We then fit the “single-site” model to these results (Figure 1B). We obtained *K_D_ =* 24.9 ± 0.7 *nM* and Δ*H*° = −11.6 ± 0.0 *kcal* · *mol*^−1^. We then repeated this analysis using *pytc*. We read the raw experimental and blank heats, uncorrected for dilution, and then globally fit *pytc’s* blank and single-site model to the two experiments using the maxiumum likelihood fitter. We used *pytc’s* built in bootstrap method to estimated parameter uncertainty by creating 1,000 psuedoreplicate datasets sampled from the heat for each shot with a standard deviation of 0.1 *kcal* · *mol*^−1^. We obtained results identical to the *Origin* results within uncertainty: *K_D_* = 24.7 ± 0.0 *nM* and Δ*H*° = −11.7 ± 0.1 *kcal* · mol^−1^ (Figure 1C). We repeated this analysis for 8 more *Ca*^2+^: *EDTA* binding experiments in three different buffers (HEPES, imidazole, and Tris). In all cases, we obtained identical results within error to those obtained by *Origin* (Figure 1D).

**Figure 1.**
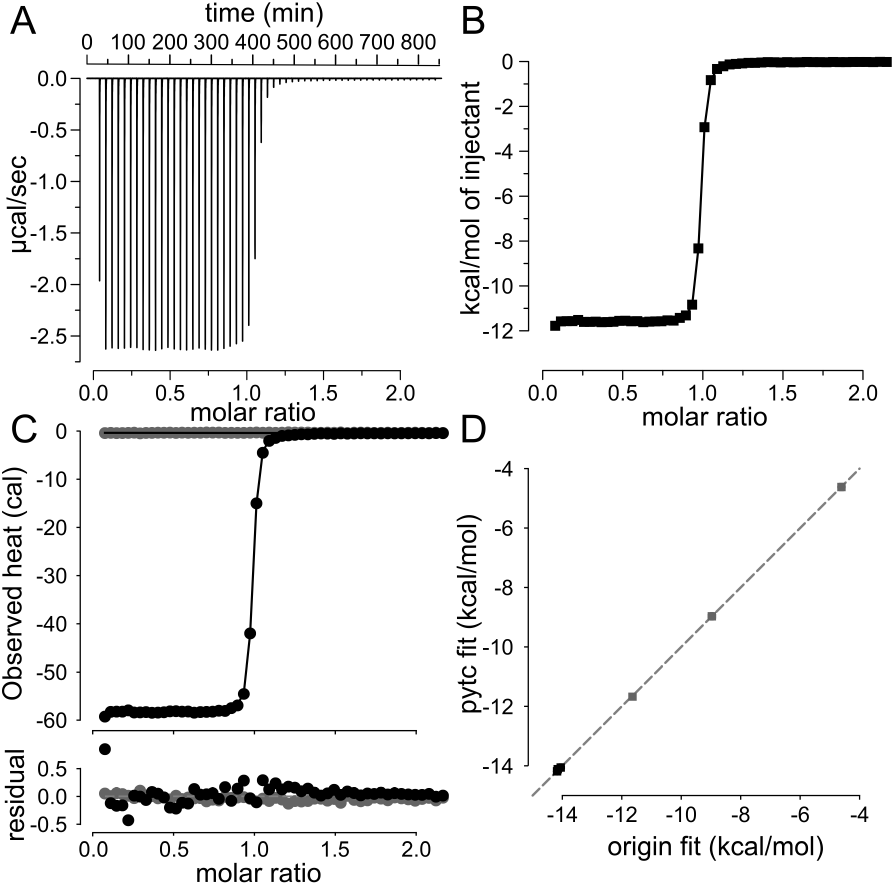
Single-site fits in *pytc* match those in *Origin*. A) Raw power curve for the binding of *Ca*^2+^ to *EDTA* (25 *mM Tris*, 100 *mM NaCl*, *pH* 7.4, 25 °*C*). B) Fit of a single-site binding model to integrated, blanked heats using *Origin* 7.0552. C) Fit of the same model with *pytc* 1.0. Dilution is accounted for in the fit model and then constrained by globally fitting the experiment (black) and blank (gray) titrations. D) Comparison of binding enthalpies (blue) and free energies (black) for *Ca*^2+^: *EDTA* interactions extracted by *Origin* and *pytc*. Plot shows 9 experiments—with three replicates each—in Tris, HEPES, and imidazole.

### Global fit to extract number of protons released

We next validated the global fitting capability of *pytc* by estimating the buffer-independent binding enthalpy 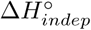 and the number of protons released or taken up on binding (*n_H_*) for the *Ca*^2+^: *EDTA* interaction. We measured binding in three buffers with different ionization enthalpies: 4.875 *kcal* · *mol*^−1^ (HEPES), 8.757 *kcal* · *mol*^−1^ (imidazole), and 11.341 *kcal* · *mol*^−1^ (Tris) (21). We used the `global_connectors.num_protons` model— which fits *K_A_*, 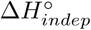, and *n_H_* —to nine experiments. This global fit yielded *n_H_* = −1.09 ± 0.02 and 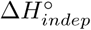 = 0.66 ± 0.13 *kcal* · *mol*^−1^ (Figure 2A). These values are in close agreement with the previously reported proton release and binding enthalpy for the *Ca*^2+^: *EDTA* interaction (22). We also performed a manual fit, plotting the measured binding enthalpies for individual experiments against the ionization enthalpy of the buffers used (3). By fitting a line to these data, we were able to extract the number of protons released *n_H_* = —1.10 ± 0.01 and the buffer-independent binding enthalpy of 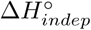 = 0.73 ± 0.10 *kcal* · *mol*^−1^ (Figure 2B). These results were indistinguishable from the global fit within the bootstrap uncertainty.

**Figure 2.**
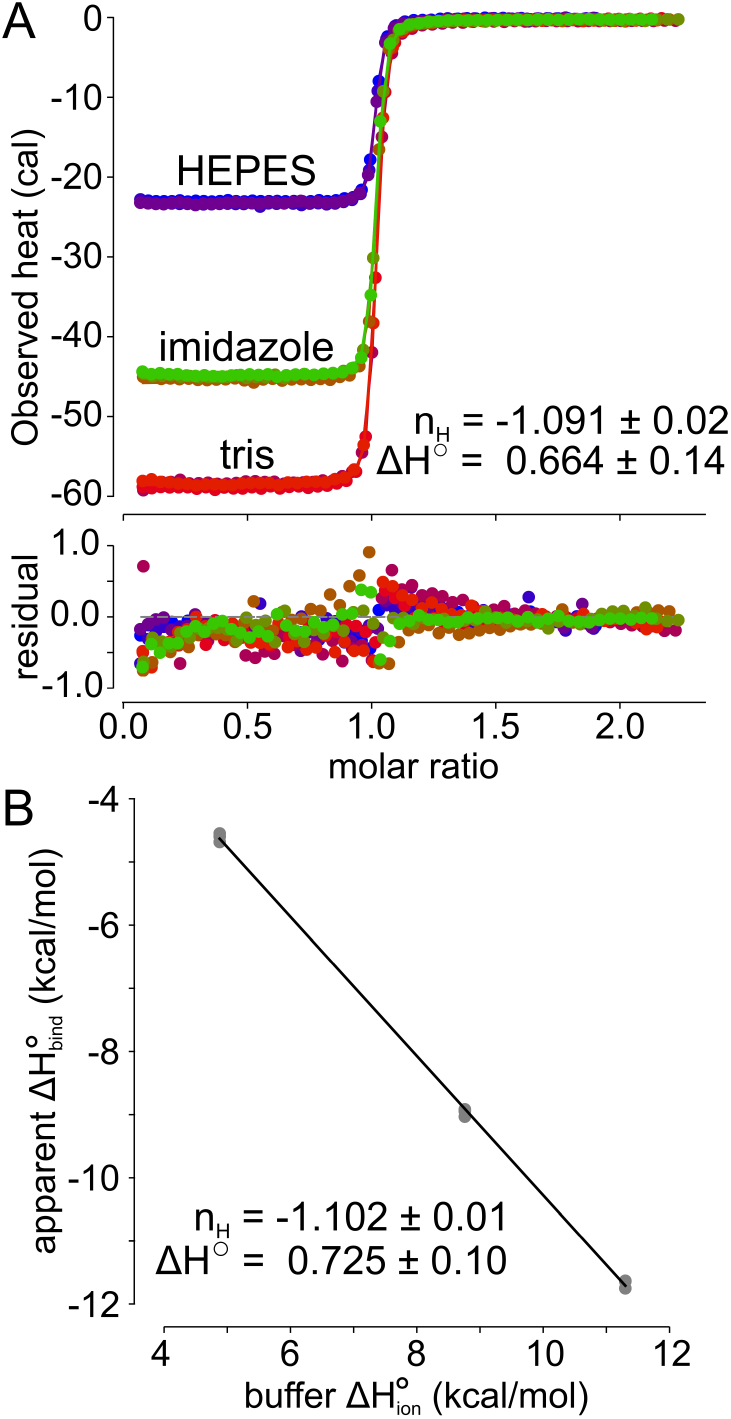
A global fit reliably extracts proton release on *Ca*^2+^: *EDTA* complex formation. A) Global fit of a proton binding model to nine individual titration experiments done in three different buffers (indicated on the graph). Colors denote experiment. We allowed the fraction competent to bind to float for each experiment, which ranged from 0.95 to 1.05. The molar ratio shown is corrected for fraction competent. B) Fit of a proton-binding model to enthalpies extracted from individual experiments versus buffer ionization enthalpy. This yields the same result as the global fit.

### Fitting multi-site models

To demonstrate the capabilities of the Bayesian global analyses we used data from a previous publication (23): the binding of *Ca*^2+^ to human S100A5. This presents a particularly challenging fitting problem. *Ca*^2+^ binds to the protein with 2:1 stoichiometry. One site is high affinity, the other low affinity. Each has its own binding enthalpy. This leads to a complex binding isotherm that is very difficult to fit in *Origin*. Further, it is difficult to resolve the entire curve in a single ITC titration, making extraction of parameters for the higher-affinity site particularly difficult. To circumvent this issue, we measured the binding of *Ca*^2+^ to S100A5 using four different titrant/titrate ratios (8×, 10×, 15×, and 18×) while keeping the protein concentration fixed across experiments (Fig 3). We performed these experiments at 25 °*C* on our ITC-200. All buffers were treated with CHELEX (≈ 1 *g* · *L*^−1^, stirring for 30 minutes) and filtered at 0.22*μm*. Prior to being loaded into the instrument, samples were centrifuged at 18,000 × *g* for 35 minutes at the experimental temperature. Shot size was 0.5 *μL* for the first shot, followed by 2.5 *μL* for all following shots. Reference power was 3 *μcal* · *s*^−1^, with stir speed set at 750 rpm.

**Figure 3.**
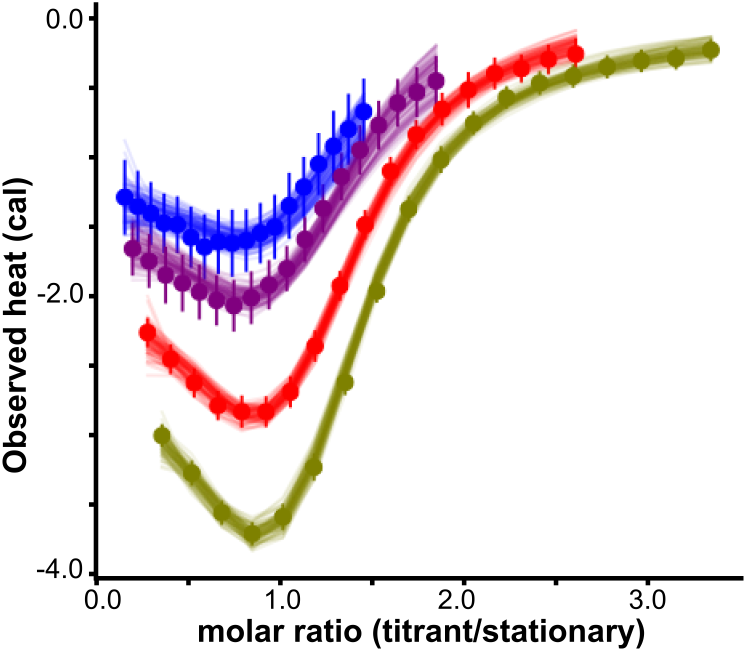
*pytc* allows global analyses of complex binding curves. Curves show titration of *Ca*^2+^ onto human S100A5 at four different titrant/titrate ratios: 8× (blue), 10× (purple), 15× (red), and 18× (green). We reported these experiments previously (23). We extracted thermodynamic parameters for a two-site binding polynomial using the Bayesian MCMC sampler in *pytc*. Points are integrated heats with uncertainty calculated using NITPIC (16). Lines are 100 curves drawn from the Bayesian posterior distribution. Simultaneous analysis of the four titrant concentrations allowed us to resolve the entire binding curve.

We then used pytc’<u>s</u> Bayesian MCMC sampler to estimate the thermodynamic parameters of a two-site binding polynomial using all four titration datasets simultaneously. We constrained the dilution heat to be between −3.0–0.0 kcal/mol and the dilution intercept between 0—10,000 kcal/mol/shot for all curves. We used uniform prior probabilities for all other parameters, which were restricted against all data simultaneously. We used the maximum likelihood estimate as a starting point, explored the likelihood surface with 100 walkers taking 20,000 steps each, and discarded the first 10 of steps as burn-in.

This strategy allowed us to resolve the entire curve simultaneously by effectively stitching together the curves from the four titrant concentrations. We were able to measure binding affinities for the high-affinity site (*K_d_*(*μM*): 0.1 ≤ 0.5 ≤ 2.7) and the low-affinity site (*K_d_*(*μM*): 1.9 ≤ 6.3 ≤ 35), which were consistent with literature values (24).

### Global Van’t Hoff analysis

To demonstrate the ability to combine the Bayesian sampler with a more complex fitting model, we performed a global Van’t Hoff analysis of a different metal-protein interaction. Previously, we discovered that the protein human S100A14 is capable of binding *Zn*^2+^ ions with relatively low affinity (*K_D_* = 50 *μM* at 25 °C) and heats (25). Here, we used *pytc* to conduct a temperature-dependent thermodynamic analysis of the binding interaction between recombinantly expressed human S100A14 and *Zn*^2+^. We titrated freshly prepared 2 *mM ZnCl_2_* onto 110 *μM* hS100A14 at 278, 283, 290, 298, 303, and 208 K using our ITC-200. Experiments were done in 25 *mM Tris*, 100 *mM NaCl* buffer at *pH* 7.4 using the ITC protocol previously described (25). We manually integrated the power curves using *Origin*.

We globally fit two variants of the Van’t Hoff model to all six datasets simultaneously using the Bayesian MCMC sampler in pytc: 1) a standard Van’t Hoff model with a 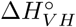 parameter and without Δ*C_p_*, and 2) an extended Van’t Hoff model with both 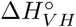 and Δ*C_p_* parameters (Figure 4A). We set the uncertainty of each integrated heat to be ±0.1 *kcal* · *mol*^−1^. We linked the 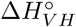, and Δ*C*_p_ parameters in a global fit to all experiments simultaneously. Other parameters, such as dilution heats, were allowed to float for each dataset separately. We used the maximum likelihood estimate as a starting guess and then explored the likelihood surface with 100 MCMC walkers, each taking 5,000 steps. We discarded the first 10 of steps as burn-in. These fits yielded global values for the 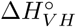 and Δ*C_p_* parameters.

**Figure 4.**
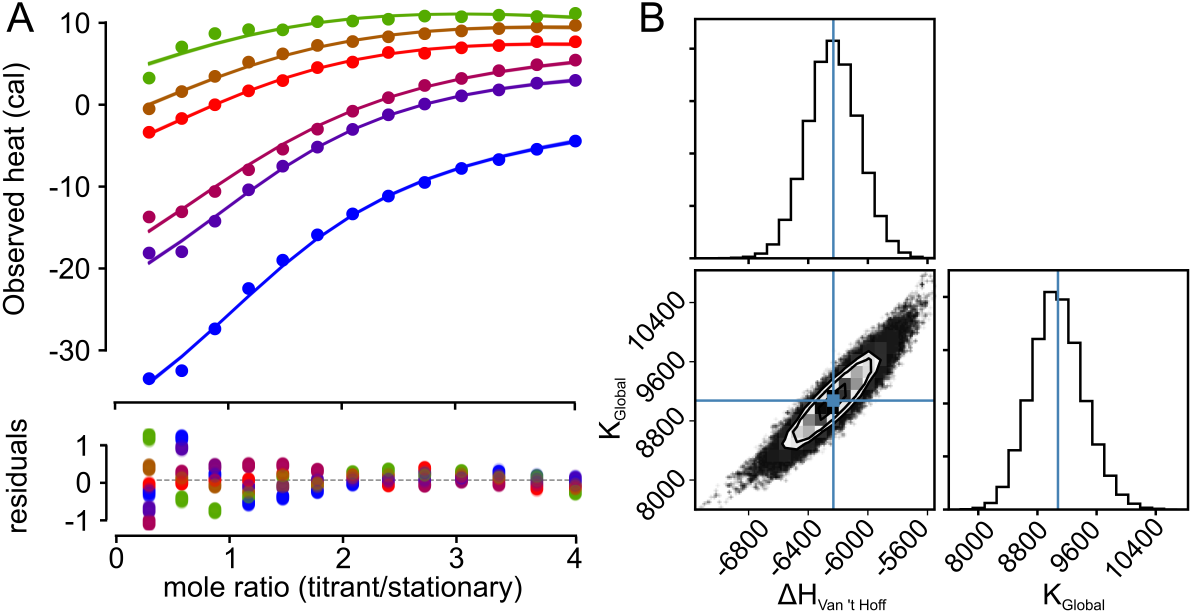
*pytc* allows for robust, global Van’t Hoff analyses. Binding of *Zn*^2+^ ions to human S100A14 was measured using ITC. Data were collected at 278, 283, 290, 298, 303, and 208 K. A) We extracted thermodynamic parameters for a single-site Van’t Hoff binding model using the Bayesian MCMC sampler in *pytc*. Points are integrated heats extracted using *Origin*. Lines are 100 curves drawn from the Bayesian posterior distribution. B) Corner plot showing correlations between the posterior distributions for the two global parameters 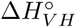 and *K_global_*. Surface was generated by 100 walkers, each taking 5,000 steps.

An additional benefit of *pytc* is the ability to easily perform model selection using the `pytc.util.compare_models` method. We used the Akaike information criterion (AIC) to assess support for the standard Van’t Hoff model relative to the extended Van’t Hoff model, finding that the standard model is strongly supported (*P* = 9.4 *x* 10^−12^). This result indicates that there is no evidence for a Δ*C_p_* associated with the binding of *Zn*^2+^ ions.

The global Bayesian fit with the standard Van’t Hoff model yielded an estimate of 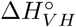 with 95 credibility region of −6.6 ≤ −6.2 ≤ −5.9 *kcal* · *mol*^−1^. Likewise, the fit returns a global estimate of the binding constant with a 95 credibility region of 1.4 × 10^4^≤ 1.8 × 10^4^ ≤ 2.2 × 10^4^ *M*^−1^, which corresponds to a maximum likelihood estimate for the *K_D_* of 56 *μM*. The Bayesian posterior surface for these parameters is shown as a corner plot (easily output using `pytc.corner_plot`()) in Fig 4B. This Van’t Hoff analysis shows the power of the Bayesian MCMC analysis and also illustrates the utility of the *pytc* API for selecting between thermodynamic models.

## Conclusion

We anticipate that *pytc* will prove useful in a wide variety of ITC analyses. Our implementation of Bayesian Markov-Chain Monte Carlo sampling, incorporation of shot-specific heat uncertainty in the fits, and inclusion of sophisticated functions for model selection make *pytc* a powerful and versatile platform for analysis of ITC data. Further, we strongly support modification and extension of *pytc* by the scientific community. Because the code is released into the public domain and has an exposed, well-documented API, we look forward to the many improvements and extensions of the code that will naturally occur as users define new models and contribute to the code base.

## Acknowledgments

The authors thank the members of the Harms lab for feedback and software testing.

## Notes

The authors declare no competing financial interests.

